# Genome-wide comparison of DNA methylation between life cycle stages of *Drosophila melanogaster* using high-throughput sequencing techniques

**DOI:** 10.1101/2020.01.06.895920

**Authors:** Saniya Deshmukh, Varada Abhyankar, Shamsudheen Karuthedath Vellarikkal, Sridhar Sivasubbu, Vinod Scaria, Deepti Deobagkar

## Abstract

*Drosophila melanogaster* undergoes holometabolous development, has very low levels of DNA methylation, and is known to possess a single known methyltransferase, dDNMT2. This study compares the DNA methylation patterns between the two life cycle stages of *D. melanogaster* using a combination of DNA immunoprecipitation and high throughput sequencing techniques.

Our results indicate, a change in the chromosomal distribution of the sparse DNA methylation concerning genes and natural transposable elements between in the embryo and the adult stages of *D. melanogaster*. The differentially methylated regions localised on genes involved in the regulation of cell cycle processes of mitotic cell divisions and chromosomal segregation. dDNMT2 knockout flies exhibited altered patterns of DNA methylation. The observed differences in DNA methylation were in genes involved in cellular communication and cytoskeletal functions. The variation in DNA methylation between the two life cycle stages is indicative that it could have a role in regulatory processes during development and, dDNMT2 may have a role as a co-factor for the hitherto undiscovered DNA methyltransferase in *D. melanogaster*.

## INTRODUCTION

DNA methylation in *Drosophila melanogaster* was first reported in the embryos by a combination of photoacoustic spectroscopy and polyclonal antibodies to be 0.008 mol percent of 5-methylcytosine (5mC) ^1,2^. These results were later corroborated by enzymatic digestion of DNA with McrBC, high-performance liquid chromatography (HPLC) and 2D-Thin layer chromatography approximately to report the presence of one 5mC molecule in 1,000-2,000 cytosines molecules in adult fly ^3^. There was no sequence-specific data in *Drosophila* genome until the analysis of the early embryos followed by sequencing of select bisulfite modified regions which revealed methylation to be predominant in non-CpG dinucleotides ^4^.

Subsequently, high-throughput sequencing of sodium bisulfite-treated immunoprecipitated fragments bearing methylation from stage 5 embryos with 10,000X coverage reported less than 1% of genome-wide DNA methylation in the stage-5 embryos ^5^. DNA samples of *D. melanogaster* in a mixed population of adult males and females (w^1118^ strain) was subjected to liquid chromatography selective reaction monitoring (LC-SRM) method to report 0.034% of methylated cytosines in the genome which is 10–100 fold below the suggested detection limit of bisulphite sequencing ^6^ whereas the female adult (Oregon-R strain) was reported to harbour 0.002% methylation of cytosines ^7^. The whole genome bisulfite-sequencing also correlated with the 55% increment in DNA methylation of the infected testes; however, this did not show any correlation with cytoplasmic incompatibility ^8^. The changes in dietary specification also do not cause any alteration in the DNA methylation of adult flies ^9^. The ultrahigh-performance liquid chromatography (UHPLC) results from twelve related species of genus *Drosophila* revealed the presence of variable DNA methylation in the genomic DNA ranging from 0.001 %5mC/C in *D. melanogaster* and 0.009 %5mC/C *D. persimilis*. The levels of 5mC change during the holometabolous development of *D. melanogaster* with 0.031 %5mC/C in the embryonic stages and 0.001 %5mC/C in the adult ^10^.

*Drosophila* belongs to the group of DNMT2-only systems which are characterised by the absence of DNMT1 and DNMT3 molecules. There has been much debate on the active DNA methyltransferase in *Drosophila*; it was suggested that the *Drosophila* DNMT2 (dDNMT2) is directly responsible for DNA methylation in *D. melanogaster* ^11^. Meanwhile, other investigations for DNMT2 substrate preference revealed that it uses RNA as a substrate and a DNMT2-like RNA methyltransferase is proposed to be a precursor for the eukaryotic DNA methyltransferases ^12,13^. It has been suggested that DNMT2 to possesses a dual specificity for RNA and DNA substrates ^14^. The dDNMT2 gene has been implied to be involved in the regulation of stress and immune response ^15–17^. Involvement of Mt2 in contributing to *Drosophila* immunity against bacterial infection in an age-dependent manner accompanied by changes in the sphingolipid metabolism has been demonstrated ^18^. The embryos of DNMT2 knockout flies have shown the persistence of DNA methylation albeit with an altered pattern clearly indicating a possibility of existence an unknown DNA methyltransferase in *Drosophila* ^5^.

The current study examines the DNA methylation patterns in the embryonic and adult stage of *D. melanogaster* by employing methylated DNA immunoprecipitation sequencing (MeDIP-Seq) technique. Presence of 5-methyl cytosine (5mC) could be detected at a very low level in the embryonic and adult stage of fruit fly. It is interesting to note that DNMT2 mutant adult flies bear DNA methylation with an altered pattern. Additionally, analysis of previously published independent whole genome bisulfite sequencing data validated the results obtained using MeDIP sequencing data and inferred the locations of DNA methylation in both embryonic as well as adult stage of *D. melanogaster*.

## MATERIALS & METHODS

### Sample preparation and DNA isolation

*D. melanogaster* late embryonic stages [Stages 12-16/8-14 hours after egg laying (AEL)] and adult males (1-3 day old) were collected from the laboratory culture of Oregon-R strain maintained on corn-meal medium under standard conditions. These samples were washed in 70% ethanol and 1X PBS; then processed for DNA isolation. Briefly, the samples were homogenised after snap-chilling in liquid nitrogen and incubated at 65 °C with RNase A (12091-021) for 4 hours; followed by phenol-chloroform extraction. Treatment with 1:2.5 parts of 5M Potassium Acetate and 6M Lithium Chloride to remove any traces of RNA was given followed by precipitation by isopropanol. The DNA samples were dissolved in TE buffer.

### Methyl-DNA immunoprecipitation sequencing (MeDIP-Seq)

The protocol by Weber & Dirk was followed with modifications ^19^. Briefly, the genomic DNA from each of the above mentioned samples was sonicated to obtain random fragments of size 100 to 500 bp. According to the manufacturer recommended protocol, fragments were end-repaired, phosphorylated and A-tailed. This DNA was ligated with Illumina adapters to the fragments. ∼4-6 µg of adaptor-ligated DNA was used for subsequent MeDIP enrichment. A 10% fraction of this adaptor-ligated DNA was set aside as input or control for each sample. A commercial monoclonal antibody against 5-methylcytidine from Eurogentec was used for immunoprecipitation. The fastq files were the input and enriched samples were obtained.

The quality assessment and trimming of the fastq file was performed by FastQC (http://www.bioinformatics.babraham.ac.uk/projects/fastqc) and Trimmomatic programs ^20^. Bowtie2 was used to align the trimmed files with the reference genome of *D. melanogaster* Release 6 (dm6) ^21^. The peak calling for the aligned files was performed using MACS version 2.1 ^22^. The MACS2 outputs for the samples were uploaded on the UCSC genome browser (https://genome.ucsc.edu/) for visualisation against dm6 reference genome.

### Whole genome bisulfite sequencing (WGBS) analysis

The available WGBS datasets 7 day old fully fed females (SRP073522), Stage 5 (2-3 hours AEL) embryo (SRR389219), bisulfite sequencing 66 regions in two samples stage 5 (SRR1191286) and MT2 knockout (SRR1191487) from NCBI-SRA repository were analysed by Bismark software version 19.0 ^23^.

### Data analysis

#### 1. Differential methylated regions (DMR) estimation

The DMR for both the MeDIP-seq and <u>WGBS</u> data were identified using suitable packages from R suite with appropriate statistical parameters. The MeDIP-seq datasets were analysed using the MEDIPS package using p-value set at 1 * 10^−5 24^. The BS-seq data analysed for DMR was processed by the methylkit package ^25^, the q-value cutoff was set at 0.1 and the minimum difference in the methylation levels was set to 5 percent value. The methylkit package was also used to map chromosome-wise DMR distribution and the associated feature annotation in the WGBS datasets.

#### 2. Gene Ontology (GO) analysis

The GO analysis for the methylated regions was performed using Cytoscape 3.5 (www.cytoscape.org/) with application ClueGO version 2.5 ^26^ and PANTHER GO ^27^.

## RESULTS

### DNA methylation & MeDIP-sequencing

In this study, the sequencing of DNA fragments enriched using an antibody against 5mC was employed to identify the differences in the patterns and associated features of the methylated sequences in the embryonic and the adult stages of *D. melanogaster*. The fold enrichment in methylated fragments in embryo stage varied between 2-5 fold whereas a 2-3 fold enrichment was observed in adult stage. The embryo and adult stages had a characteristic and typical distribution of DNA methylation with respect to the number of genomic fragments and their chromosome-wise distribution [Figure 1]. Also, relatively higher DNA methylation in embryo was evident by presence of a greater number of methylated regions in embryo as compared to adult stage. The maximum number of methylated regions was found on chromosome 3L in the embryo and chromosome 2R of the adult stage. Y chromosome had similar number of methylated regions in both embryo and adult stages of *D. melanogaster* [Figure 1a]. The length of chromosome fractioned out in the immunoprecipitation to the total length of individual chromosomes was also variable in the two developmental stages of *D. melanogaster* [Figure 1b]. The total length of the individual chromosome had no effect on the pull-down rate by the antibody in both the samples suggesting a stage-specific distribution of 5mC in localised regions. The feature annotation of the immunoprecipitated fragments revealed that both geneic [Figure 1c] and transposable elements (TE) were methylated [Figure 1d].

**Figure 1:**
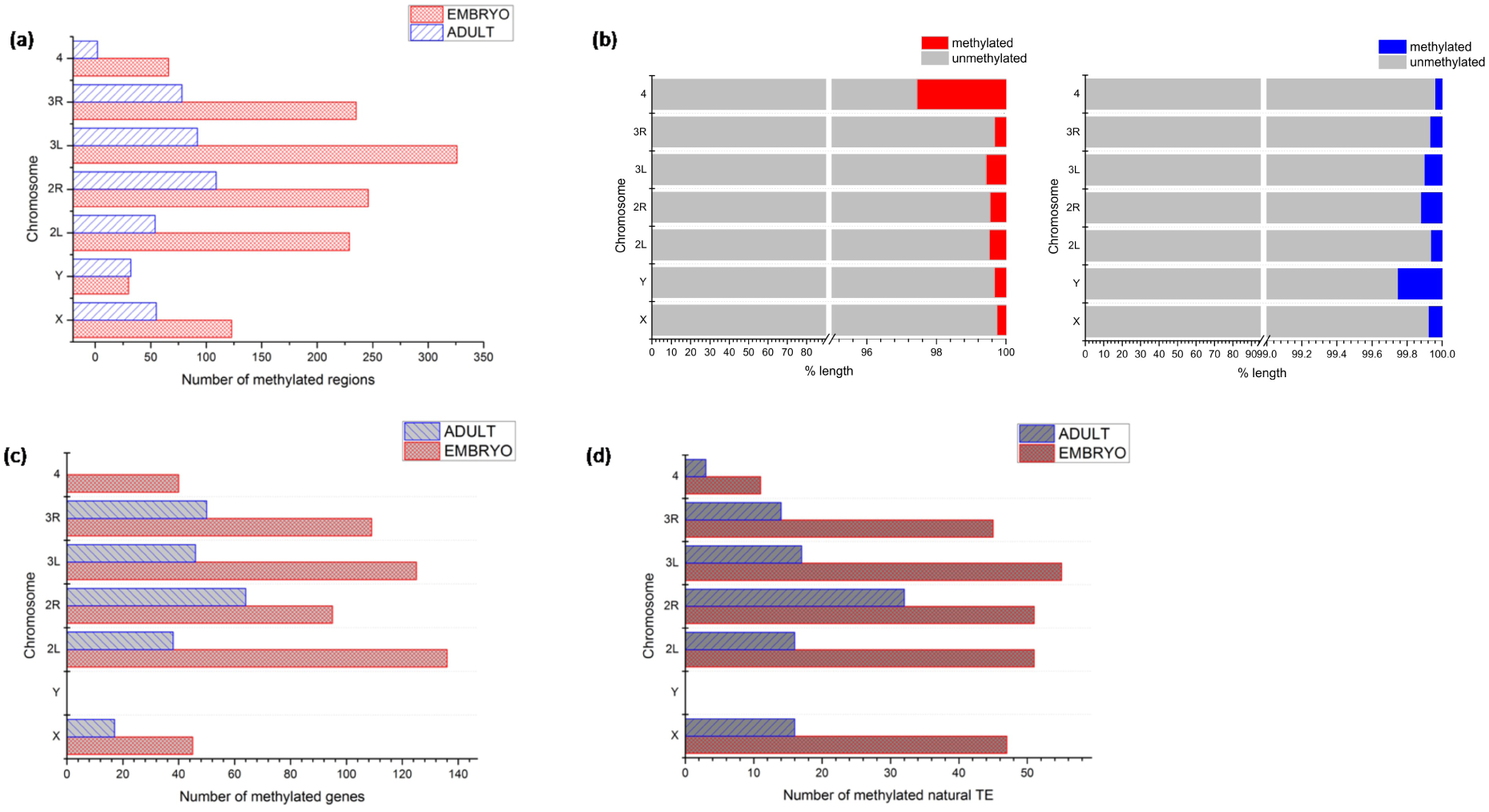
The representation of chromosomal distribution of DNA methylation in the embryo and adult stages of *D. melanogaster*. (a) the number of immunoprecipitated fragments per chromosome (b) the length (in percent) of each chromosome in the pulled-down fraction as compared to the total chromosomal length in embryo (red) and adult (blue) samples (c) the number of annotated genes present in the methylated regions per chromosome (d) the natural transposable elements methylated per chromosome.

The embryo stage had double the number of methylated genes as compared to the adult stage. The maximum number of gene-bearing regions was located on chromosome 2L in embryo while on chromosome 2R in the adult. Although Y chromosome has methylation, no methylated genes were identified on the Y chromosome in both the developmental stages. Additionally, in adult flies, chromosome 4 lacked the presence of methylated genes. Transposable elements (TE) in the genome were also found to be more in the embryonic stage than the adult stage. The maximum TEs were located chromosome 3L and chromosome 2R in the embryo and in the adult respectively. Interestingly, in the adult stage of *D. melanogaster* number of methylated regions found on chromosome 4 showed maximum overlap with the methylated transposable elements.

When the gene ontology of methylated genes in the embryo was analysed; genes involved in biological processes such as like sister chromatid separation, intra and inter cellular signalling, cell cycle regulation which are essential for the embryo development were identified. The biological processes associated with methylated genes are diverse as the developing embryo has multiple signalling and developmental pathways that are actively working on the transition of an embryo to the subsequent life cycle stage (supplementary figure 3a). On the contrary, in the adult, the methylated genes are associated with more stringent set of biological process linked with the higher-order brain functions like short-term memory formation, axon genesis and cell cycle regulation (supplementary figure 3b).

The DREME analysis of ±50 bp region around the summit of the methylated peak regions in the two stages confirmed DNA methylation in *Drosophila* is present in short, specific motifs. These methylated regions are enriched in low complexity motifs, particularly CT-rich in the embryo while they are CC-rich in the adult stage. This is indicative of an asymmetric and rare pattern in the genomic methylation (supplementary figure 4).

Our analysis has identified substantial differences in the pattern of DNA methylation between the embryonic and adult stages of development. Since, the dDNMT2 molecule is the only known methyltransferase in *D. melanogaster.* The distribution of DNA methylation in wild-type (Mt2^+/+^) and the mutant of dDNMT2 (Mt2^-/-^) in the adult stage was compared by employing the immunoprecipitation in order to understand the effect of the absence of dDNMT2. The dDNMT2 mutant adult flies harboured DNA methylation in low levels and there were striking differences between the wild-type (Mt2^+/+^) when compared with Mt2 mutant (Mt2^-/-^) [Figure 2]. The total immunoprecipated regions obtained corresponding to each chromosome are different in case of the wild-type and mutant adults. More methylated regions were identified on chromosome 2R of the wild-type while in Mt2 adult mutant, chromosome 3R demonstrated the presence of the highest number of methylated regions [Figure 2a]. Chromosome 4 had the least methylated regions in both the datasets while chromosomes X and Y were comparable in wild and mutant datasets. It was observed that a greater percentage of chromosomes 3R and Y in Mt2 mutant flies was immunoprecipitated than wild-type [Figure 2b]. The chromosome 2R bore maximum genes [Figure 2c] and TE [Figure 2d] in the wild type. In case of the mutant adults, however, the maximum number of methylated genes and TEs are present on chromosomes X and 3L respectively.

**Figure 2:**
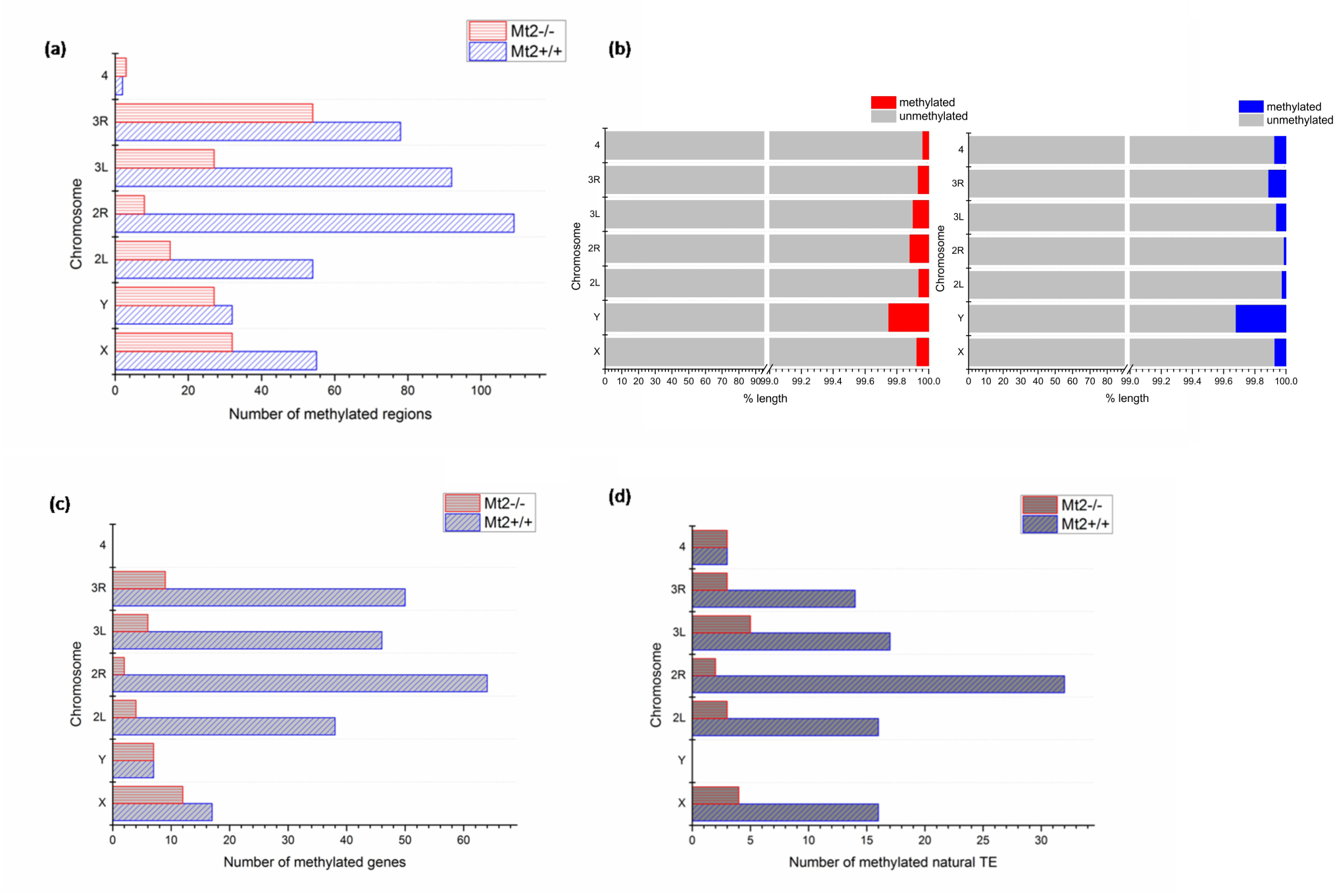
The representation of chromosomal distribution of DNA methylation in the wild-type (Mt2^+/+^) and Mt2 mutant (Mt2^-/-^) genotypes of *D. melanogaster*. (a) the number of immunoprecipitated fragments per chromosome (b) the length (in percent) of each chromosome in the pulled-down fraction as compared to the total chromosomal length in Mt2^+/+^ (red) and Mt2^-/-^ (blue) samples (c) the number of annotated genes present in the methylated regions per chromosome (d) the natural transposable elements methylated per chromosome.

The DREME analysis of methylated regions in the wild-type and Mt2 mutant revealed the presence of distinct and specific pattern in the methylated sequences. Unlike the wild-type, the DNA motifs from the mutant sample are guanine-rich implying a shift in the DNA methylation due to absence of dDNMT2 (supplementary figure 4).

### Differentially methylated regions in embryo and adult stages

Gene ontology analysis of the methylated regions in the embryo and adult stages identified genes involved in functions related to chromosomal segregation and mitotic divisions. For example, in the embryo and adult stage, DMR like rough deal (rod) and non-claret disjunctional (ncd) are involved in the kinetochore-microtubule assembly and mitotic progression are differentially methylated (Table 1).

**Table 1:** GO processes in DMR between developmental stages and Mt2 mutant (Mt2^-/-^)

| Gene ontology of DMR late embryos and adult male |  |  |
| --- | --- | --- |
| GO Identifier | GO Term | Term P value |
| GO:0010941 | regulation of cell death | 0.002 |
| GO:0007059 | chromosome segregation | 0.001 |
| GO:0098813 | nuclear chromosome segregation | 0.001 |
| GO:0000819 | sister chromatid segregation | 0.000 |
| GO:0007067 | mitotic nuclear division | 0.002 |
| GO:0000070 | mitotic sister chromatid segregation | 0.000 |
| Gene ontology of DMR wild-type (Mt2<sup>+/+</sup>) and Mt2 mutant (Mt2<sup>-/-</sup>) |  |  |
| GO Identifier | GO Term | Term P value |
| GO:0030594 | neurotransmitter receptor activity | 0.001 |
| GO:0005230 | extracellular ligand-gated ion channel activity | 0.001 |
| GO:0034330 | cell junction organization | 0.001 |
| GO:0016337 | single organismal cell-cell adhesion | 0.002 |
| GO:0032970 | regulation of actin filament-based process | 0.001 |
| GO:0032956 | regulation of actin cytoskeleton organization | 0.001 |

In the absence of dDNMT2, the mutant adult possesses fewer methylated regions than in the wild-type adult stage. The DMR are involved in functions related to cytoskeletal organisation and inter-cellular signalling processes (Table 1). The GO of methylated genes for the MeDIP datasets are represented in supplementary data (supplementary figure 3).

### DNA methylation & whole genome bisulfite sequencing (WGBS)

In order to establish patterns and locations of DMRs using an alternative technique, the available WGBS datasets from independent studies from public repositories for the embryo and adult stage were analysed. As described in the materials and methods, the embryonic stage data was obtained from a study on the stage 5 embryos ^5^ and the adult stage data from adult whole body sample ^8^. This analysis reiterated that embryo stage had higher and consistent DNA methylation than the adult stage [Figure 3a and 3b]. Predominant DNA methylation was seen in the CHH nucleotide context in the embryo followed by CpG and CHG context while adult had a very low level of methylation; it was 0.5% in the CHH trinucleotide sequence and 0.4% in CHG and CpG context (Table 2). In stage 5 embryos, the frequency of occurrence of sites with 100% DNA methylation is about 20 whereas the whole body adult sample exhibited a low frequency of occurrence of methylation and the possibility of any given site being methylated is less than 10 per cent [Figure 3a and Figure 3b]. The chromosomes X, 2R and 3R in the adult had relatively higher methylation as compared to the stage 5 embryo in the CHH context [Figure 3c] and the differential DNA methylation was predominantly in the intergenic regions and not the promoters [Figure 3d]. The DNA methylation of the Ubx locus was analysed as a case study and both samples showed the presence of variably methylated cytosines in the adult sample as compared to fewer methylated cytosines at distinct locations in the stage 5 embryo [Figure 3e]. The remaining DNA methylation was seen in the CHG and CpG context. In the adult samples, the chromosomes X and 3R harboured more methylation as compared to the stage 5 embryos in the CHG context and a predominant difference in the DNA methylation of promoter regions of embryo and adult was evident (Supplementary figure 5.1). The chromosomes 2L and 3R in the adult samples are the only chromosomes to bear decreased methylation (statistically significant) as compared to the stage 5 embryo in the CpG context; there is a predominant difference in the DNA methylation in the exon regions of embryo and adult sample (Supplementary figure 5.2).

**Figure 3:**
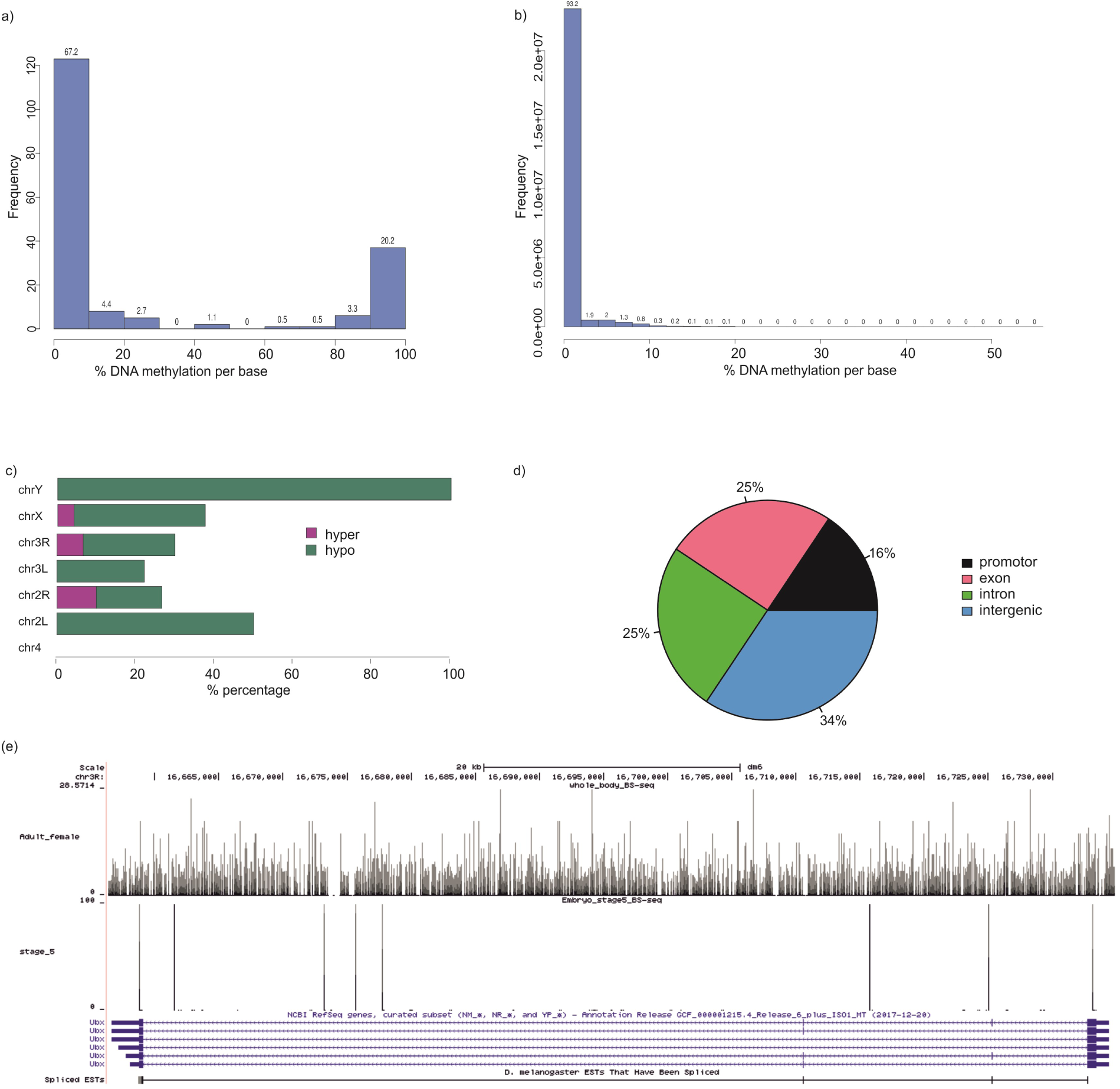
The comparison of DNA methylation in CHH context by BS-seq in developmental stages of *D. melanogaster*. (a) descriptive statistics on percent DNA methylation in the stage 5 embryo sample (b) descriptive statistics on percent DNA methylation in the adult sample (c) chromosomal distribution of differentially methylated regions between the embryo and adult sample (d) annotation of the differentially methylated regions (e) UCSC browser representation of a methylated region bearing the gene region Ubx.

**Table 2:** Distribution of DNA methylation between the embryonic and adult samples.

| DNA cytosine methylation between stages of development |  |  |
| --- | --- | --- |
|  | Adult whole body | Stage 5 embryo |
| C methylated in CpG context | 0.40% | 3.90% |
| C methylated in CHG context | 0.40% | 3.30% |
| C methylated in CHH context | 0.50% | 5.20% |

The bisulfite-sequencing data for stage 5 embryo with respect to wild-type (Mt2^+/+^) and Mt2 mutant (Mt2^-/-^) genotypes in the public repositories is available for 66 regions ^5^ was examined for patterns in sequence context. Methylation was present in the CHH context in both genotypes (Supplementary 6.1 [a] & [b]). Differentially methylated regions were identified in the Mt2^-/-^ sample against the Mt2^+/+^ sample. The chromosomes X, 2L, 2R, 3L and 3R in the Mt2^-/-^samples showed relatively more methylation as compared to the Mt2^+/+^ in the CHH context (Supplementary figure 6.1 c). The locations of methylation with respect to specific cytosines were different at some loci of the representative analysis of DNA methylation of the kirre locus showed the presence of an additional peak in Mt2^-/-^ sample as compared to the Mt2^+/+^ sample (Supplementary figure 6.1 e). The wild-type and mutant displayed methylation features with differences in the introns with respect to the CHH, CHG and CpG context (Supplementary figures 6.1, 6.2 & 6.3).

### Comparison of DNA methylation from MeDIP-seq and WGBS

Although, MeDIP-seq can identify the presence of DNA methylation in a genomic region it does not give information about the exact location of 5mC in this sequence. In order to assess the DNA methylation pattern, in the embryonic and adult stages of *D. melanogaster* using alternative method, we have analysed and compared WGBS data from public repositories for the embryo and the adult stages. The methylated genes from the MeDIP-seq dataset common with the genes from WGBS data were identified using the data from UCSC genomes [Figure 4].

**Figure 4:**
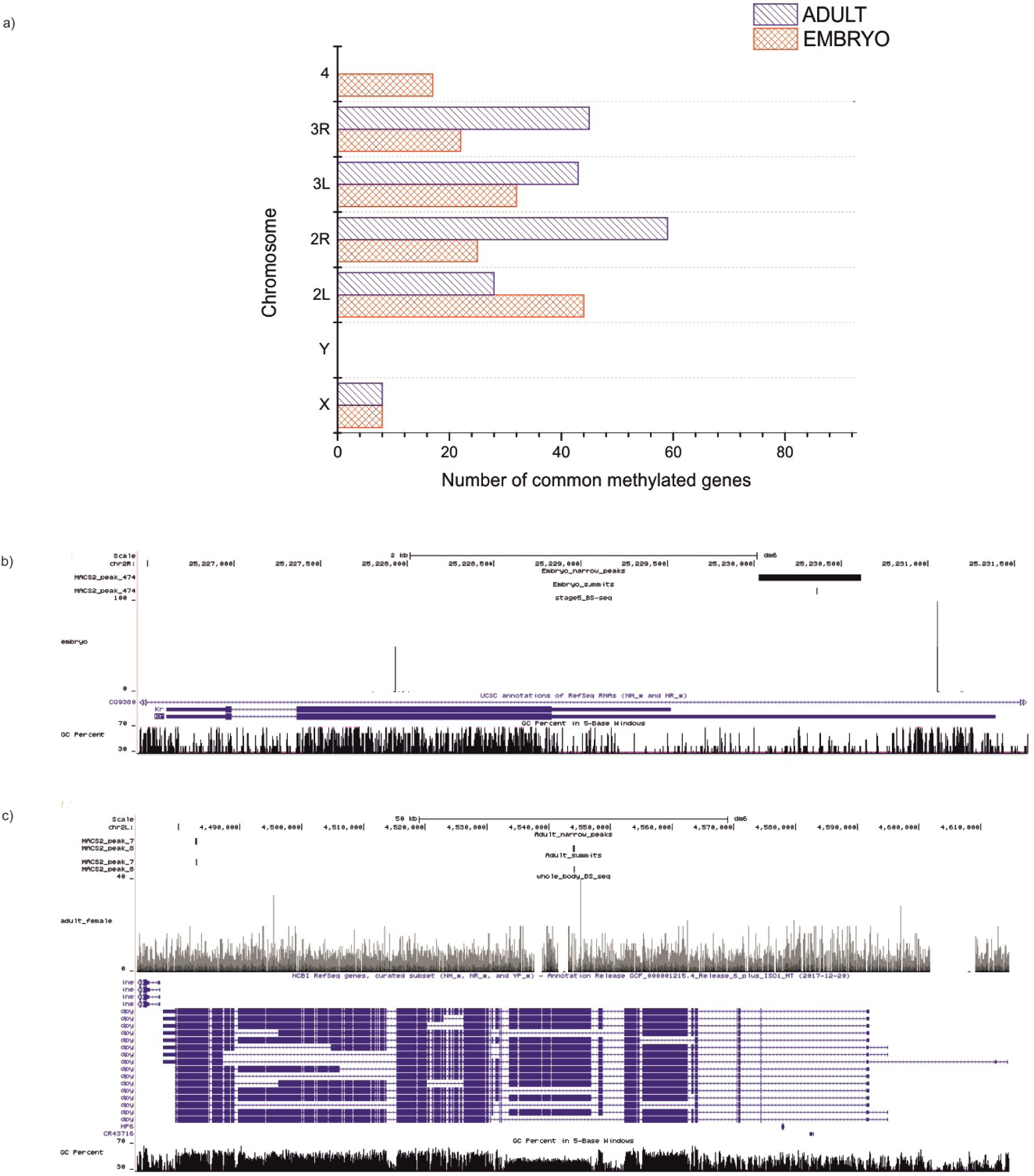
The comparison of MeDIP seq data with the available WGBS data. (a) the number of methylated genes from each stage identified by the MeDIP-seq found common with the WGBS data using UCSC. The graphical representation of the methylated genes detected by MeDIP-seq and BS-seq techniques(b) Kruppel [Kr] in the embryonic stages (c) dumpy [dpy] in the adult stage. The top tracks represent the MeDIP-seq data for both genes with the peak regions and the peak summits for the anti-5mC antibody interacting sites. The second panel depicts data analysed from an independent WGBS studies on stage 5 embryos and adult females for the same genes; the vertical bars mark the location of the methylated cytosine in the gene. The other UCSC tracks besides the sequenced datasets are GC content and Refseq genes.

In the embryo stage, the genes bearing methylated regions between the two datasets were compared; all methylated genes from the MeDIP-seq identified using the UCSC refseq data were found to be present as a subset of the WGBS dataset. However, the methylated peaks from the MeDIP-seq experiment from stages 12-16 did not share any direct overlap with the methylated loci identified from the WGBS of stage 5. For example, chromosome X had 8 methylated genes from the MeDIP-seq dataset of late embryos which were all a subset of the WGBS data from stage 5. When the MeDIP-seq data for adult sample (1-2day old males) and WGBS data (9-10 day old females) was analysed, several common regions could be identified as methylated with a direct overlap of most peaks. Thus, both approaches could identify some common methylated regions and the difference in the methylation pattern could be explained by the differences in the age and sex of the sample used. As demonstrated [Figure 4a], the chromosome X harbours 17 methylated genes from MeDIP-seq of adult male and 8 genes from this dataset bear methylated loci identified from the WGBS of whole body of adult female. The methylated genes are visualised, Kruppel from the embryo [Figure 4b] and dumpy from the adult [Figure 4c] by both the techniques.

## DISCUSSION

Although, the presence of DNA methylation and the role of DNMT2 as a DNA methyltransferase has been a debatable issue in *D. melanogaster* ^11,28,29^, recent developments in the field have established presence of low levels and asymmetric pattern of DNA methylation in *D. melanogaster* ^5,30^. It has been documented that the levels of DNA methylation in *Drosophila* with change respect to developmental stages ^10,31^, the comprehensive analysis regarding the location and chromosomal distribution was not available. The current study provides a comparative analysis of DNA methylation from embryo and adult stages by MeDIP-seq and WGBS. This revealed DNA methylation in the embryo stages is higher and has a different chromosomal distribution in comparison to the adult stage indicating the dynamic change in levels and distribution of 5mC during development. There was no correlation between DNA methylation and the expressed sequence tags (EST) data in both the life cycle stages of *D. melanogaster*.

The possible involvement of DNA methylation in the regulatory pathways of *Drosophila* can be better understood by additional exploration of the positioning and interaction of 5mC with other epigenetic modulators or DNA-binding proteins. Our results indicate that DNA methylation of genes involved in mitotic progression is most altered between the embryo to the adult stage. The DMR between these stages are located in the intergenic and promoter regions in the non-CpG context whereas with respect to the CpG context the difference is predominant in the exon regions. The functional importance of DNA methylation in controlling neural aspects like behaviour, social interactions, caste determination, and memory are better understood in the eusocial systems like honey bees and wasps which possess an epigenetic machinery similar to mammals ^32^. This indicated that DNA methylation could be associated with functions like parent-of-origin marking of epigenome. DNA methylation in carpenter ants is associated with histone dependant the spatial organization of chromatin within active genes ^33^. Whether the low level of DNA methylation in *D. melanogaster* is involved in epigenome marking for trans-generational transfer of information, regulation of gene expression is open to speculation. These findings among eusocial taxa are widely studied in association with CpG content ^34^. This proves to be a shortcoming in extrapolating the role of DNA methylation with sparse and predominantly non-CpG based DNA methylation like *D. melanogaster*. However, in this study we have examined the patterns and location of 5mC with respect to *Drosophila* development. From the current analysis carried it can be postulated that *Drosophila* DNA methylation appears to regulate specific cellular processes as compared to the range of functionality in other systems. The possibility of other epigenetic modifications (miRNA, histone modifications etc.) taking over regulatory roles assigned to DNA methylation in the well-known developmental processes cannot be ruled out.

dDNMT2 is the only known methyltransferase in *D. melanogaster* and its role in tRNA methylation has been elaborated in literature ^14^. Earlier report ^5^ demonstrated presence of DNA methylation with an altered patterns in the Dnmt2 mutant embryos. In the current study, Dnmt2 mutant adult (Figure 2) exhibited methylation at a fewer sites with an altered pattern. A shift in the DNA methylation of the DMRs in the intron regions was observed in wild-type (Mt2^+/+^) and Mt2 mutant (Mt2^-/-^) of stage 5 embryos (supplementary figure 6). This thus, raises question about the ability of Dnmt2 to methylate DNA, as its loss neither completely erased the DNA methylation nor retained same pattern of methylation. Hence, it can be postulated that a novel methyltransferase might be working closely with Dnmt2 to establish the methylation pattern observed in wild-type flies. Dnmt2 mutant flies were shown to be susceptible temperature, oxidative and viral stress ^16,17,29^; additionally, the ability to handle bacterial infection in these flies is reported to reduce in an age dependent manner accompanied by changes in hemocyte shape and lipid profile ^18^. In the light of the presented data, further analysis of the DNA methylation in the adult Dnmt2 mutant flies might reveal its role in handling biotic or abiotic stress.

MeDIP-seq and WGBS techniques have certain specific advantages in the analysis of DNA methylation profiles. MeDIP-seq cannot identify the exact location of 5mC in the genome an advantage offered by WGBS studies. Nonetheless, WGBS has its own shortcomings when detecting very low levels of DNA methylation ^6^. Our analysis has revealed that the narrow peaks obtained in the MeDIP-seq data represented the region of DNA which interacted with the antibody against 5mC and the summit is the nucleotide which showed maximum interaction with the antibody. In majority of the peaks analysed in the present study the summit corresponds to a C or G indicating methylation in the forward or reverse strand respectively (supplementary figure 1 & 2). Despite the observed overlap between results from the two techniques, differences were observed in the exact location of the DNA methylation between samples. This can be attributed to the low propensity of per base methylation in *Drosophila* and difference in the age and sex of sample used for the BS-seq and MeDIP-seq from the two stages.

In conclusion, there is a differential pattern of DNA methylation observed in the development of *D. melanogaster* and the adult has much lower and variable pattern of methylation. However, it is unlikely that this low level of DNA methylation is involved in direct regulation of gene expression via mechanisms similar to mammals or in honey bees. It appears that Dnmt2 definitely modulates or alters patterns of DNA methylation in the fruit fly. The possibility of the existence of additional methyltransferase molecule or a protein complex involved in methyltransferase activity cannot be completely eliminated as the dDNMT2 knockout had persistence of DNA methylation.

## DATA ACCESS

The high throughput data generated for this thesis has been uploaded to the NCBI-SRA data repository (https://www.ncbi.nlm.nih.gov/sra) under the bioproject identifier PRJNA431532: The analysis of methylated DNA sequences in two developmental stages of *Drosophila melanogaster* (File accession numbers: SRR6770835, SRR6770836, SRR6770834,SRR6770832, SRR6770833 and SRR6770837). SRA datasets used NCBI-SRA are SRR6770835, SRR6770834, SRR6770836, SRR6770837, SRR6770832 and SRR6770833, the studies are duly cited in text^5,8^.

## AUTHOR CONTRIBUTIONS

SD and VA designed and carried out the experiments, performed data analysis. SKV carried out the MeDIP sequencing for embryo and adult samples. SD, VA, VS, SS and DD designed and coordinated the study and drafted the manuscript.

## ACKNOWLEDGMENTS

SD is supported by UGC-UPE Phase II Biotechnology [UGC-262(A)(1)]. VA is supported by fellowship from CSIR, Govt. of India. DD is supported by UGC-UPE Phase II Biotechnology [UGC-262(A)(1)] and SPPU-DRDP grant. Mr. Rijit Jayarajan and Mr. Ankit Verma for help with sequencing experiments.

