## Supplemental data for "Genome-wide comparison of DNA methylation between life cycle stages of *Drosophila melanogaster* using high-throughput sequencing techniques"

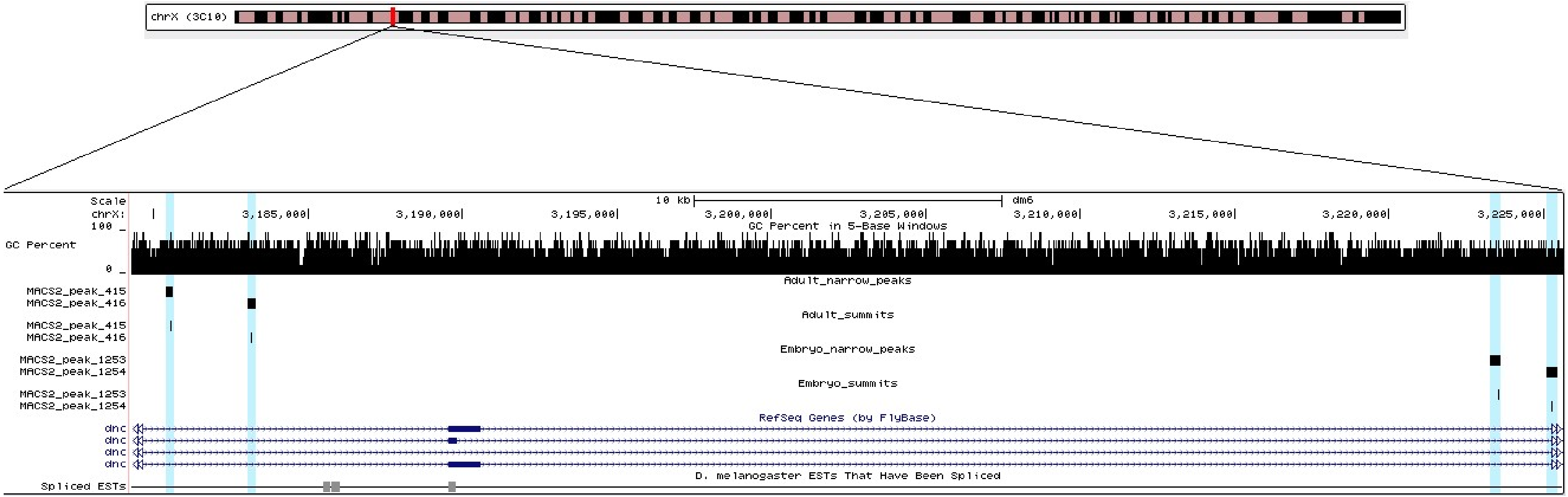


Supplementary figure 1: The methylated genes in the MeDIP-seq analysis of the embryo and adult stages of *D. melanogaster* the regions highlighted in blue are methylated the rectangular box represents the peak region and the vertical line represents the peak summit for the two samples; the other UCSC tracks besides the embryo and adult are GC content and Refseq genes for *D. melanogaster* gene dunce (dnc), a learning associated gene with behavioural parameters like learning, memory and signalling is methylated in the both the samples.

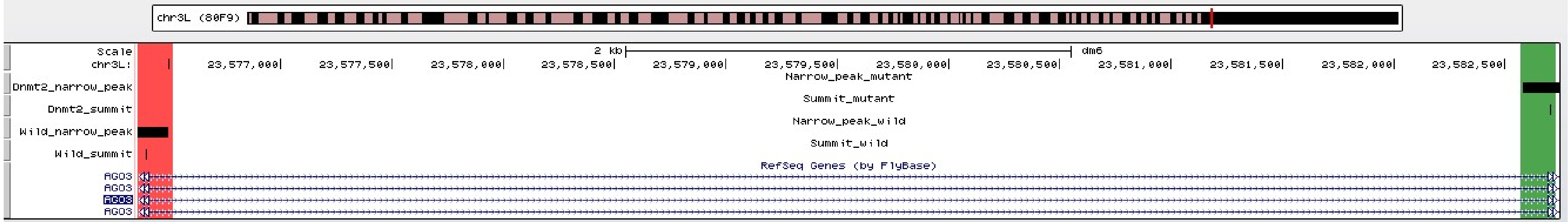


Supplementary figure 2: The methylated genes in the MeDIP-seq analysis in the wild type (Mt2$^{+/+}$) and Mt2 mutant (Mt2$^{-/-}$) genotypes of *D. melanogaster* are highlighted in red and green respectively. The other UCSC tracks besides the MeDIP-seq are GC content and Refseq genes for *D. melanogaster* gene Argonaute (Ago3), a germline-specific Piwi family protein, which interacts with Piwi-interacting RNAs (piRNAs) during germline development.

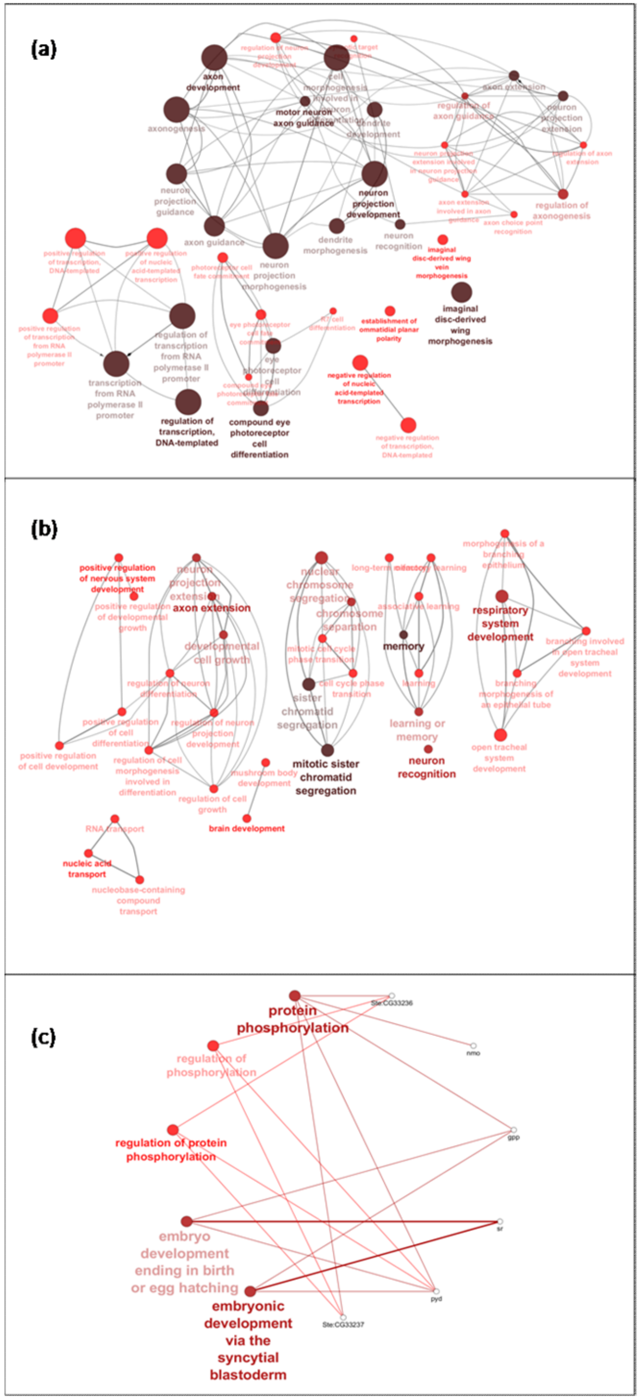


Supplementary figure 3: The gene ontology of regions methylated in MeDIP-seq samples (a) late embryo (b) the wild type ($Mt2^{+/+}$) and (c) Mt2 mutant ($Mt2^{-/-}$)

Methylated motifs
**EMBRYO**

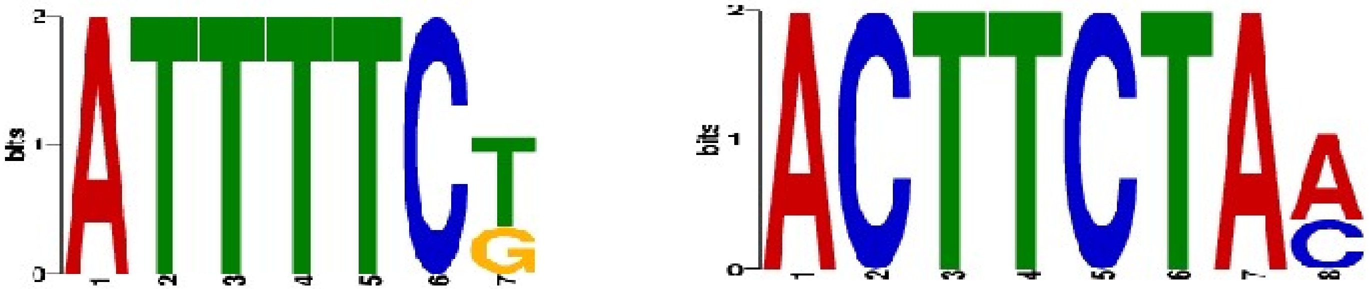


Methylated motifs
**ADULT**

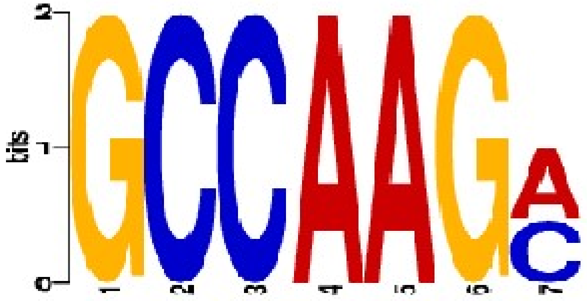


Methylated motifs
**MT2 MUTANT**

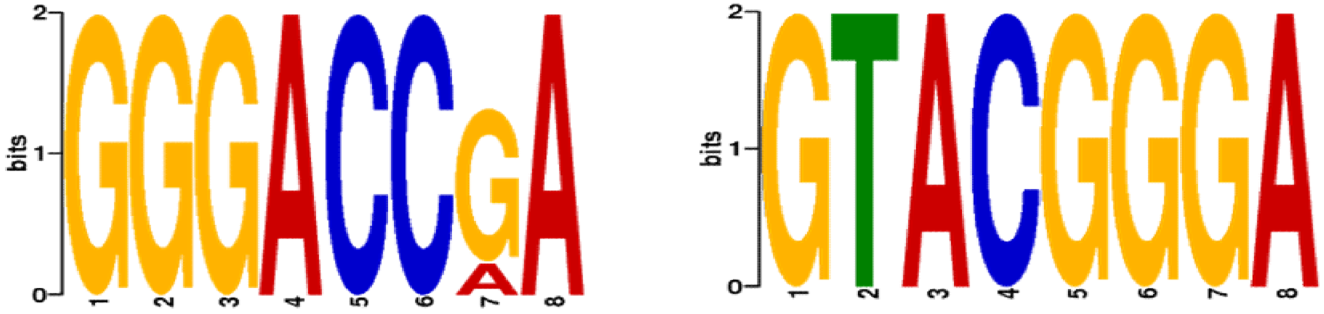


Supplementary figure 4: The DREME analysis of ±50 bp region around the summit of the methylated peak confirms DNA methylation in short and specific motif which are rare.

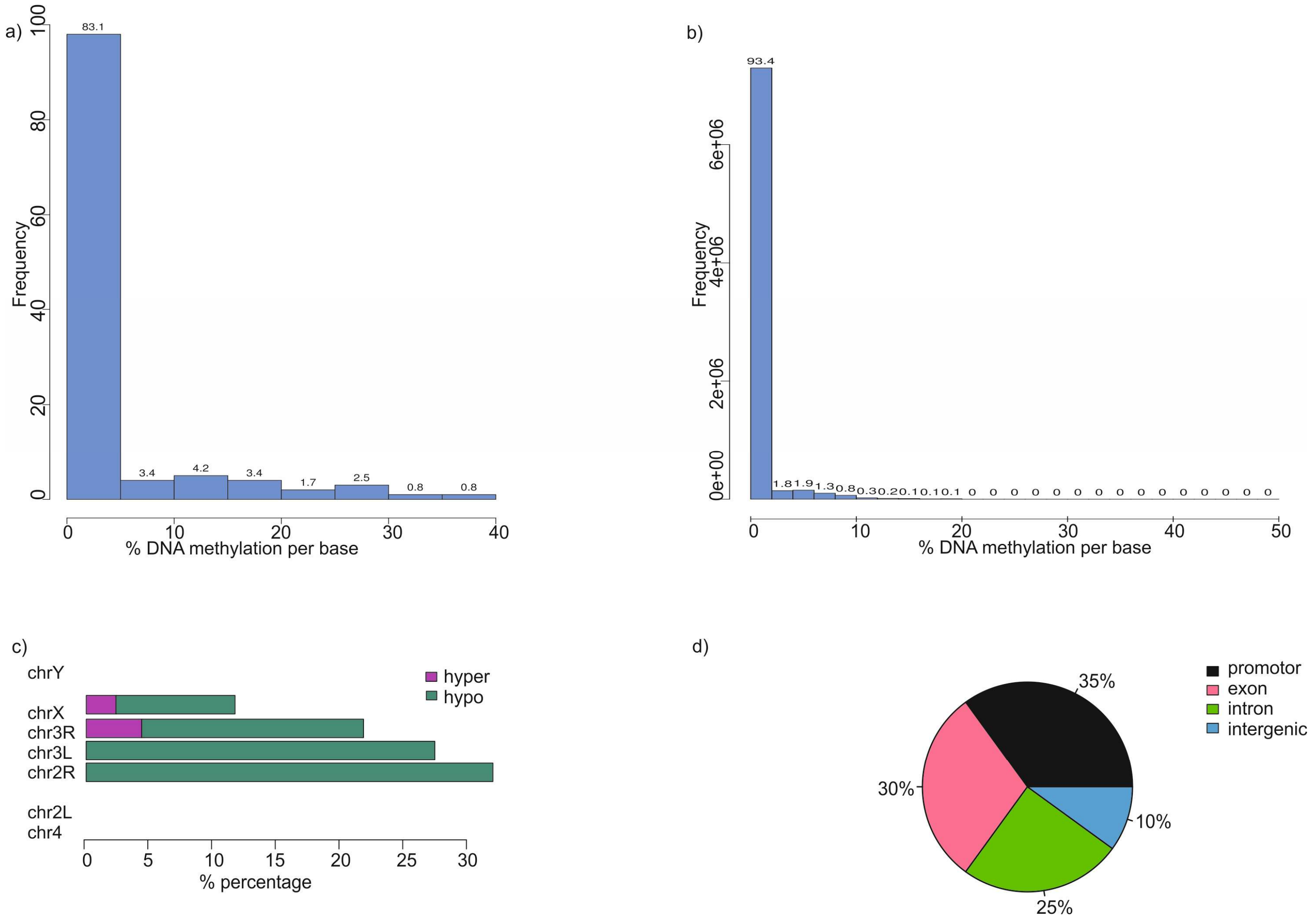


Supplementary figure 5.1: The comparison of DNA methylation in CHG context by BS-seq in developmental stages of *D. melanogaster*: (a) descriptive statistics on percent DNA methylation in the stage 5 embryo sample (b) descriptive statistics on percent DNA methylation in the adult sample (c) chromosomal distribution of differentially methylated regions between the embryo and adult sample (d) annotation of the differentially methylated regions.

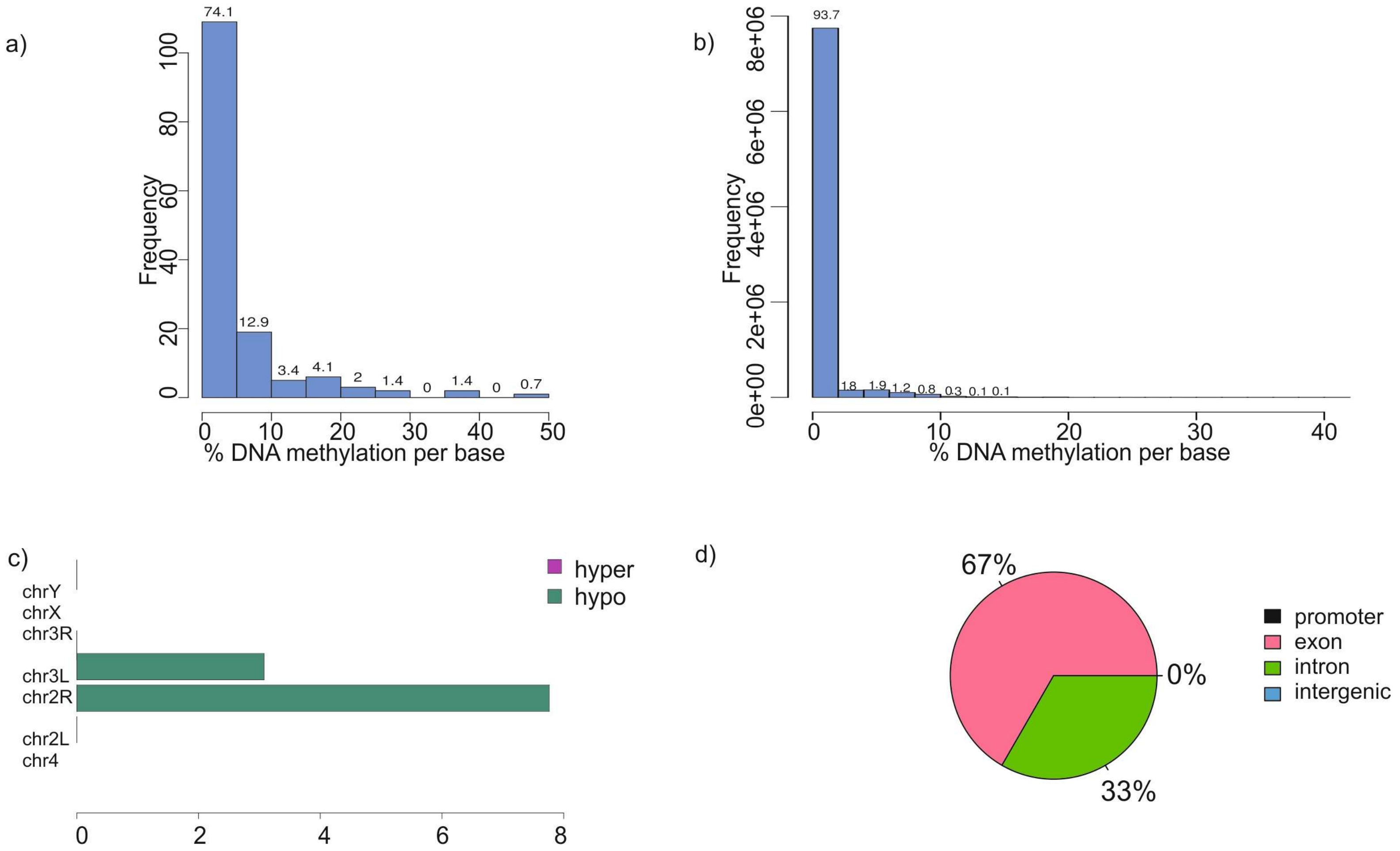


Supplementary figure 5.2: The comparison of DNA methylation in CpG context by BS-seq in developmental stages of *D. melanogaster*: (a) descriptive statistics on percent DNA methylation in the stage 5 embryo sample (b) descriptive statistics on percent DNA methylation in the adult sample (c) chromosomal distribution of differentially methylated regions between the embryo and adult sample (d) annotation of the differentially methylated regions.

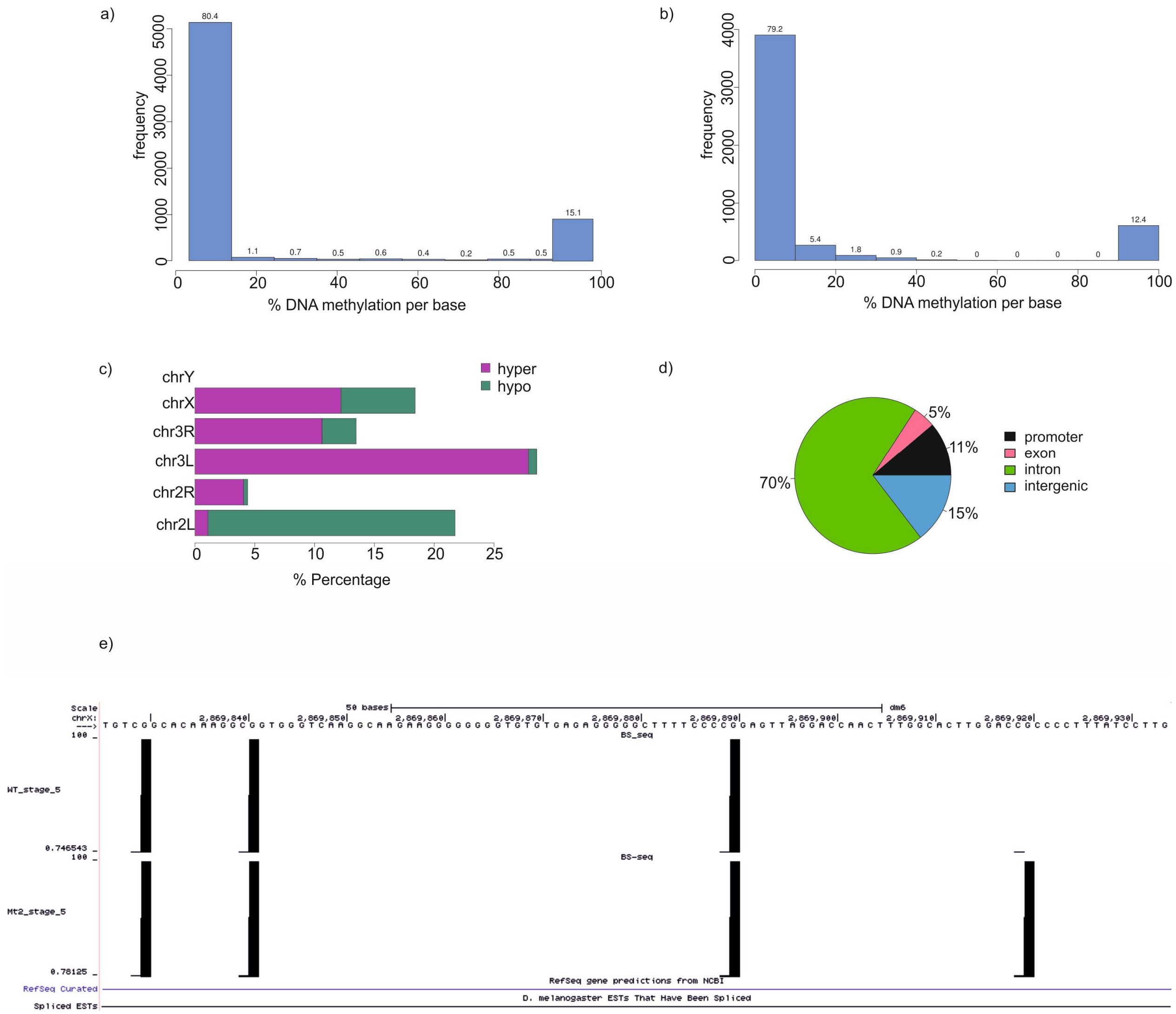


Supplementary figure 6.1: The comparison of DNA methylation in CHH context by BS-seq in stage 5 embryos of *D. melanogaster* with reference to DNMT2: (a) descriptive statistics on percent DNA methylation in the stage 5 embryo wild type ($Mt2^{+/+}$) sample (b) descriptive statistics on percent DNA methylation in the stage 5 embryo Mt2 mutant ($Mt2^{-/-}$) sample (c) chromosomal distribution of differentially methylated regions between the $Mt2^{+/+}$&$Mt2^{-/-}$ sample (d) annotation of the differentially methylated regions (e) UCSC browser representation of a methylated region bearing the gene region kirre.

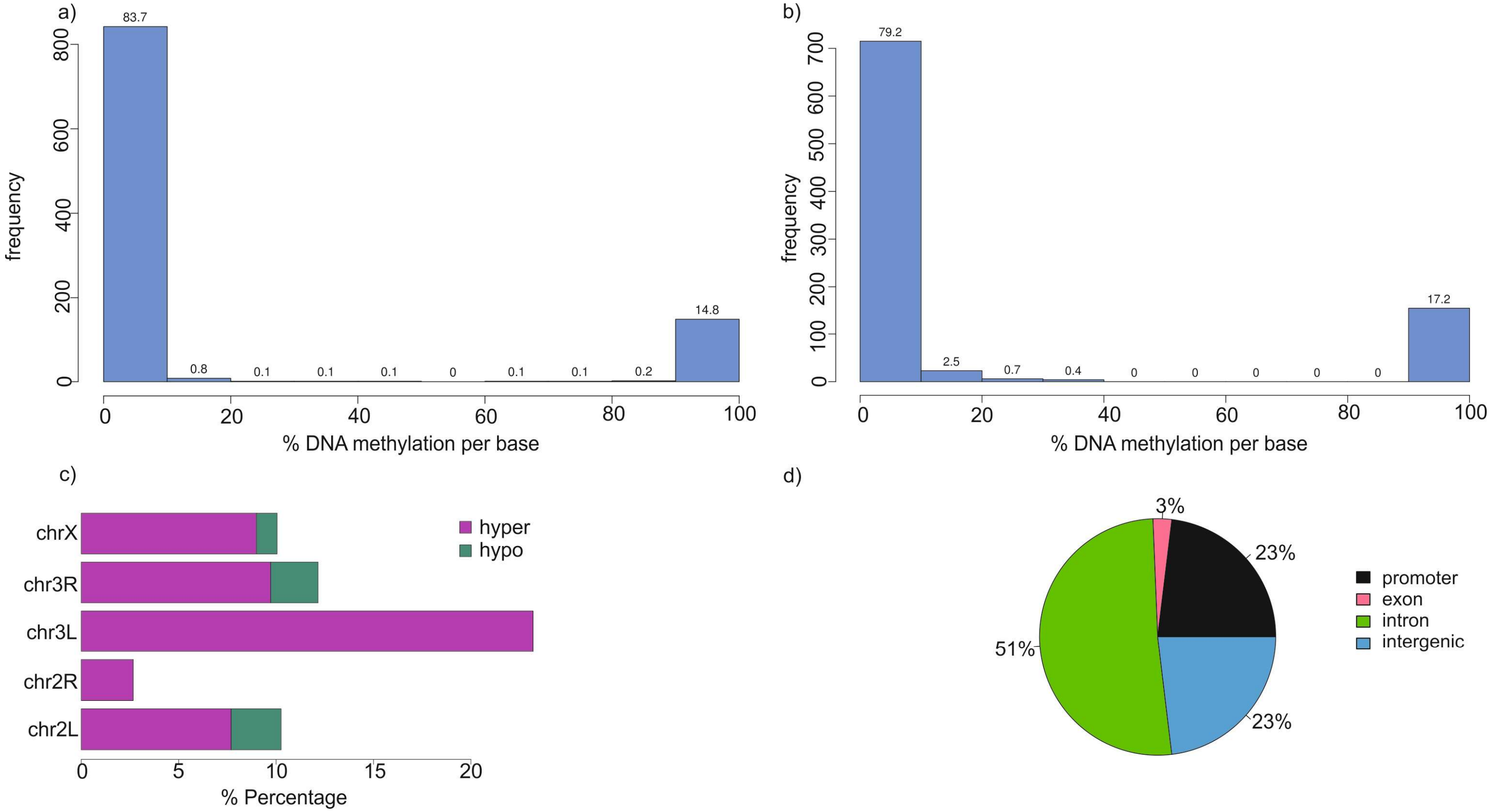


Supplementary figure 6.2: The comparison of DNA methylation in CHG context by BS-seq in stage 5 embryos of *D. melanogaster* with reference to DNMT2: (a) descriptive statistics on percent DNA methylation in the stage 5 embryo wild type ($Mt2^{+/+}$) sample (b) descriptive statistics on percent DNA methylation in the stage 5 embryo Mt2 mutant ($Mt2^{-/-}$) sample (c) chromosomal distribution of differentially methylated regions between the $Mt2^{+/+}$&$Mt2^{-/-}$ sample (d) annotation of the differentially methylated regions.

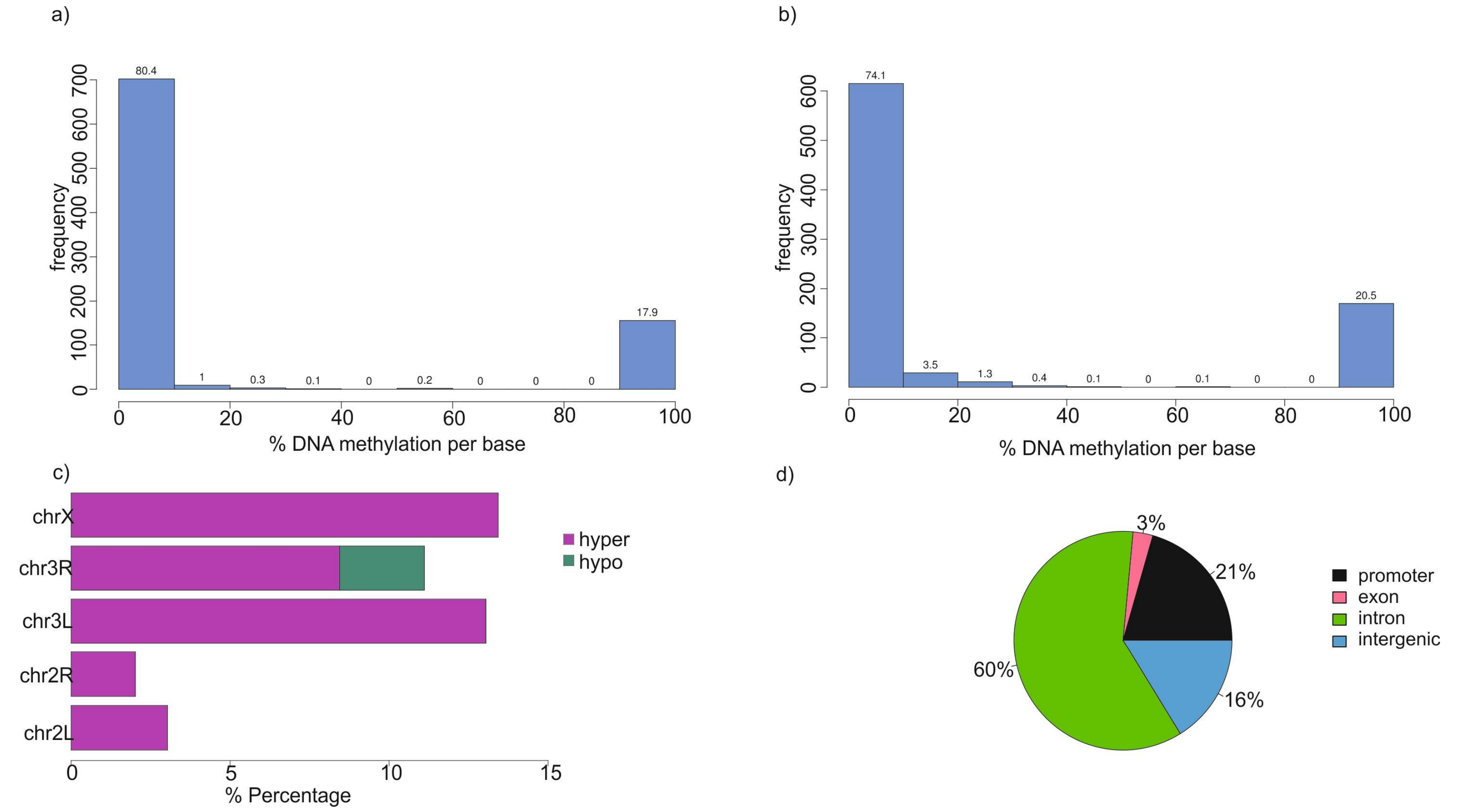


Supplementary figure 6.3: The comparison of DNA methylation in CpG context by BS-seq in stage 5 embryos of *D. melanogaster* with reference to DNMT2: (a) descriptive statistics on percent DNA methylation in the stage 5 embryo wild type ($Mt2^{+/+}$) sample (b) descriptive statistics on percent DNA methylation in the stage 5 embryo Mt2 mutant ($Mt2^{-/-}$) sample (c) chromosomal distribution of differentially methylated regions between the $Mt2^{+/+}$&$Mt2^{-/-}$ sample (d) annotation of the differentially methylated regions.
